# Enrichment of glycoRNAs using galactose oxidase, hydrazide chemistry, and glycosidase digestion

**DOI:** 10.1101/2024.06.02.597007

**Authors:** Xinyu Miao, Jianbo Deng, Xiaotong Wang, Sheng-Ye Wen, Zeyang Zhou, Shuwei Li, Xiaodong Yang, Shuang Yang

## Abstract

Ribonucleic acid (RNA), essential for protein production and immune function, undergoes glycosylation, a process that attaches carbohydrates to RNA, creating unique glycoRNAs. These sugar-coated RNA molecules regulate immune responses and may be related to immune disorders. However, studying them is challenging due to RNA’s fragility. Therefore, a robust method for identifying glycosylated RNA is important. To address this, we optimized methods for enriching and identifying glycoRNAs, opening doors to explore their potential interactions with immune receptors and tumor suppression. Our approach involved investigating factors such as preservation solutions, enzyme buffers, digestion temperature, and incubation time. We successfully achieved efficient digestion of both N-linked and O-linked glycoRNAs at room temperature using 25 mM ammonium bicarbonate, demonstrating the effectiveness of this method. Additionally, RNA preservation in RNAlater at -80°C allows controlled release of glycoRNAs within hours. While sequential digestion of different glycoRNA types is possible, significant degradation occurs after the first enzyme step. Thus, we recommend separate harvesting for each type of glycoRNA. These optimized protocols, utilizing SPCgRNA and TnORNA methods, pave the way for further research on N- and O-glycoRNAs in health and disease.

## Introduction

RNA, a remarkably versatile molecule, plays a wide range of functions within the cell. Beyond its crucial role in translating DNA’s genetic code into proteins, RNA also modifies other RNA molecules and regulates gene expression, allowing cells to adapt to different environments and developmental stages. This intricate regulation involves various chemical modifications, such as N6-methyladenosine (m^6^A) ^1^, 5- methylcytosine (m^5^C) ^2,3^. These modifications are not random; they significantly impact immune cell biology and influence diverse cellular processes, including cell death, proliferation, differentiation, and responses to DNA damage ^4^. For instance, the presence of m6A on messenger RNA (mRNA) can influence gene expression and ultimately control cell development and fate ^5^.

Recent studies have revealed that RNA can undergo post-transcriptional modifications by carbohydrates. Fialho and colleagues demonstrated that 2,4,6-triaminopyrimidine (TAP), a possible ancestral nucleobase of RNA, can be glycosylated in water by non-ribose sugars ^6^. Nucleotides within RNA can be modified at various positions (heterocyclic base, sugar, and phosphodiester linkage) by enzymes called glycoenzymes, resulting in glycosylated RNA (glycoRNA) ^7,8^. In 2021, Bertozzi and colleagues utilized click chemistry to reveal that RNA on the cellular surface can be decorated by sialoglycans ^9^. Building on previous discoveries, Ma et al. implemented a method for spatial visualization of glycoRNAs using a sialic acid aptamer and RNA in situ hybridization-mediated proximity ligation assay (ARPLA) ^10^. Zhang et al. revealed that specific glycoRNAs on the surface of neutrophils are essential for these cells to interact with and migrate through blood vessel linings, enabling them to reach inflammation sites ^11^. Flynn et al. recently identified the modified RNA base acp3U as an attachment site for N-glycans, utilizing a combination of peroidate oxidation-aldehyde ligation and mass spectrometry ^12^. With the increasing recognition of glycoRNAs’ diverse roles in health and disease, robust techniques for their identification are crucial to unlock a deeper understanding of their biological functions.

The cis-diol structure of carbohydrates is well-known to readily undergo oxidation by periodate, generating aldehydes useful for chemical conjugation ^13,14^. This approach, widely used in studying protein glycosylation, involves oxidizing glycopeptides and covalently conjugating them to a resin containing aldehyde-reactive groups ^15,16^. Sodium periodate effectively oxidizes various carbohydrates, including sialic acid, fucose, galactose, and N-acetylgalactosamine (GalNAc) ^17,18^. However, it exhibits lower reactivity towards mannose and N-acetylglucosamine (GlcNAc) ^19^. A key limitation of periodate oxidation is its lack of specificity. At high concentrations, it can damage the entire glycan structure and even nucleotides. Additionally, RNA nucleotides lack the cis-diol groups that enable periodate oxidation in carbohydrates. Sodium periodate readily oxidizes the 5’ and 3’ termini of RNA, further exacerbating its degradation by RNases ^20^. This vulnerability stems from RNA’s single-stranded nature and its ribose sugars, unlike the more stable deoxyribose sugars found in double-stranded DNA. Therefore, sodium periodate’s lack of specificity renders it unsuitable for RNA oxidation. Emerging as a more promising alternative, galactose oxidase specifically targets galactose (Gal) and N-acetylgalactosamine (GalNAc) while preserving RNA integrity ^21^. Leveraging this principle, we developed a method called solid-phase capture of oxidized N-linked glycoRNAs (SPCgRNA) for successful enrichment and identification of N-glycoRNAs ^22^.

This study aimed to optimize the enrichment and identification of glycoRNAs using the solid-phase capture of oxidized N-linked glycoRNAs (SPCgRNA) method. While our previous work successfully enriched glycoRNAs from cultured cells, tissues pose a greater challenge due to rapid degradation by endogenous RNases ^23^. To address this, we compared RNA quality and integrity isolated with and without a protective buffer. Enriched glycoRNAs were then analyzed for length and relative abundance. We further investigated key parameters for optimization: galactose oxidase (GAO) treatment duration and amount, and glycoRNA release conditions including digestion temperature, time, buffer pH, and the use of sequential glycosidase digestion. This optimization of the SPCgRNA protocol not only enables enrichment of glycoRNAs with improved quality and integrity, but these conditions can also be applied for Tn-containing O-glycoRNA (TnORNA) enrichment and identification.

## Experimental procedures

### Chemical and Reagents

All materials were purchased from Sigma Aldrich unless specified otherwise. The primary reagents are listed in **Table 1**. DEPC Water, Gel Red, 6× DNA Loading Buffer, 1 M Tris-HCl (pH 6.8), and TRIzol were ordered from Shanghai Beyotime Biotechnology Co., LTD (Shanghai, China), while RNA ladder was purchased from Abclonal (Wuhan, China). Isopropanol (IPA), trichloromethane (TCM), and ethanol were obtained from Shanghai Macklin Bio-chemical Co., Ltd. (Shanghai, China). Galactose oxidase (GAO) was ordered from Sigma-Aldrich (St. Louis, MO, USA), horseradish peroxidase (HRP) from Dalian Meilun Biotechnology Co., LTD (Dalian, China), UltraLink™ Hydrazide Resin, PNGase F and O-glycosidase from New England BioLabs (Ipswich, MA, USA), GalNAcEXO from Genovis (Cambridge, MA, USA). Ammonium bicarbonate (NH4HCO3) was obtained from Sinopharm Chemical Reagent Co., Ltd. (Shanghai, China).

**Table 1.** Reagents used in glycoRNA enrichment are listed. The initial RNA amount, along with various enzymes such as galactose oxidase (GAO), PNGase F, O-glycosidase, and hydrazide resins, are provided for each test.

| **Material** | **Concentration** | **Volume (μl)** |
| --- | --- | --- |
| RNA | / | 40 μg |
| GAO | 1/3 U/μl | 3 |
| HRP | 0.8 U/μl | 0.8 |
| DMSO | / | 17 |
| Hydrazide resin | / | 50 |
| PNGase F | 10 mg/μl | 0.6 |
| O-glycosidase | 200 U/μl | 1 |
| GalNAcEXO | 20 U/μl | 1 |

### Total RNA extraction and detection

Following ethical approval (Second Affiliated Hospital of Soochow University Ethics Committee), human colorectal tissues were collected from patients with written informed consent. As shown in **Figure 1**, the isolation process involved homogenizing 50 mg of RNAlater-stored tissue with 1 mL TRIzol using a manual homogenizer on ice to prevent RNA degradation. After a 5-minute incubation at room temperature, 0.2 mL of chloroform was added, followed by vortexing to mix. The sample was then incubated for 2-3 minutes to allow phase separation. Finally, centrifugation at 12,000 × g for 15 minutes at 4°C separated the phases. The aqueous phase, containing the RNA, was carefully transferred to a new tube, mixed with 1 mL of isopropanol (IPA), and incubated for 10 minutes. In the final centrifugation step, the RNA is precipitated at 12,000 × g for 10 minutes at 4°C.

**Figure 1.**
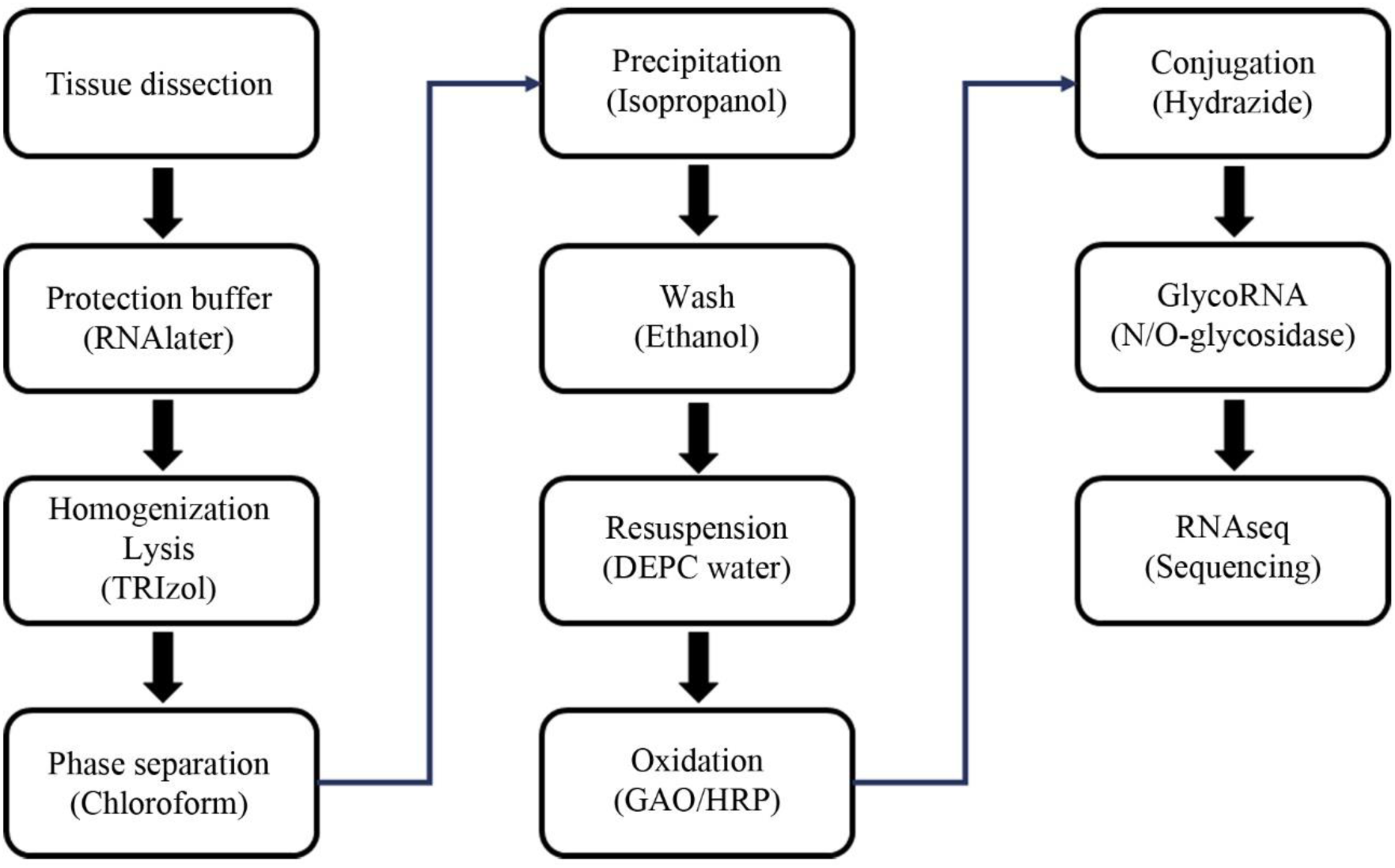
General workflow of tissue RNA extraction for glycoRNA enrichment. Tissue collection and storage: Tissue is dissected into small pieces and stored in RNAlater buffer at -80°C to preserve RNA integrity before further analysis. RNA extraction: The tissue is homogenized, and the lysate is incubated with TRIzol solution to isolate total RNA. Chloroform is then added to induce phase separation. RNA purification: The RNA-containing aqueous phase is collected, and the RNA is precipitated using isopropanol. The RNA pellet is further purified by washing with ethanol. RNA elution and oxidation: The purified RNA is eluted in DEPC water. GlycoRNAs are then oxidized using sodium periodate (NaIO4) and horseradish peroxidase (HRP). GlycoRNA enrichment: The oxidized glycoRNAs are conjugated to a hydrazide resin. The enriched glycoRNAs are subsequently released from the resin using glycosidases for RNA sequencing (RNAseq) analysis.

To remove contaminants, the RNA pellet was washed with 1 mL of 75% ethanol after removing the solution. The sample was then centrifuged at 8,000 × g for 5 minutes at 4°C. The supernatant was aspirated, leaving approximately 30 μL of solution at the bottom of the tube. This wash step was repeated for 1 minute at the same speed and temperature. The RNA pellet was then air-dried until transparent and resuspended in 30 μL of DEPC water (Beyotime, Shanghai, China). RNA purity (A260/A280 and A260/A230 ratios) and yield were measured using a Nanodrop instrument (Thermo). The integrity of the total RNA was assessed by agarose gel electrophoresis. Briefly, the samples were loaded onto a 1% agarose gel and run at 150 V for 20 minutes in 0.5× TAE buffer.

### GlycoRNA isolation and detection

Total RNA (approximately 40 μg) meeting quality requirements underwent oxidation. The sample was incubated with 1 U each of galactose oxidase (GAO) and horseradish peroxidase (HRP) in 85 μL DEPC water containing 20% (v/v) DMSO for 2 hours at 25°C. This step selectively targets glycoRNAs containing galactose (Gal), N-acetylgalactosamine (GalNAc), or fucose (Fuc). The oxidized RNA was then incubated with 50 μL of Ultralink hydrazide resin on a 1000 rpm shaker for 2-3 hours at 25°C to capture the glycoRNAs. Finally, the resin was washed five times with 200 μL of DEPC water to remove non-specifically bound RNA or other molecules.

N-glycosylated RNA cleavage from the hydrazide resin involved incubating it with 100 μL reaction buffer containing 25 mM ammonium bicarbonate (NH4HCO3) and 0.6 μL PNGase F (500 U/μL) for 3 hours at 25°C with shaking at 1000 rpm. To ensure complete cleavage, the eluate was collected, and the incubation step was repeated. The released RNA was stored at -80°C for future RNA sequencing (RNAseq) analysis. O-glycosylated RNA cleavage followed a similar protocol: the hydrazide resin was incubated with a reaction buffer containing 25 mM NH4HCO3, 1 μL O-glycosidase (200 U/μL), and 1 μL GalNAcEXO (20 U/μL) (TnORNA) for 3 hours at 25°C with 1000 rpm shaking, mirroring the N-glycosylation cleavage conditions. The eluate was collected and the incubation step repeated once to ensure complete cleavage. The released RNA samples were then stored at -80°C. It’s important to note that alternative reaction buffers, such as 25 mM Tris-HCl or PBS, can be used with this protocol. In such cases, only the buffer needs to be substituted, while the overall sample preparation procedure remains similar.

## Results and discussion

### Preserving RNA and glycoRNA integrity with stabilization buffer

To assess the effectiveness of RNA stabilization buffers on RNA yield and length, we compared tissues stored in two commercially available options: RNAlater (Thermo Fisher Scientific, Waltham, MA, USA) and RNAsafe (Nanjing Apollomics, Nanjing, Jiangsu, China). We contrasted these with unpreserved controls. Approximately 50 mg of tissue from each set was stored under two conditions: 1) in RNAlater or RNAsafe, and 2) directly in a sterilized tube without any stabilization buffer. Following extraction, all tissues were immediately stored at -80°C. **Figure 2** depicts the RNA integrity of triplicate tissue samples from three patient cohorts, after one month following surgery. Samples were either preserved in RNAlater or stored in empty tubes. RNA integrity is typically assessed based on the relative abundance of the 28S, 18S, and 5S ribosomal RNA (rRNA) bands. In high-quality RNA, the 28S and 18S bands are dominant, while severe degradation is indicated by a stronger 5S band. Our results clearly demonstrate significant RNA degradation in tissues stored without a stabilization buffer (empty tubes). Conversely, tissues stored in RNAlater exhibited minimal degradation, highlighting its effectiveness in preserving RNA integrity of human tumor tissues using this preservation buffer. Moreover, extracted total RNA from RNAlater-protected tissue yielded significantly higher glycoRNA enrichment (**Figure 2**A).

**Figure 2.**
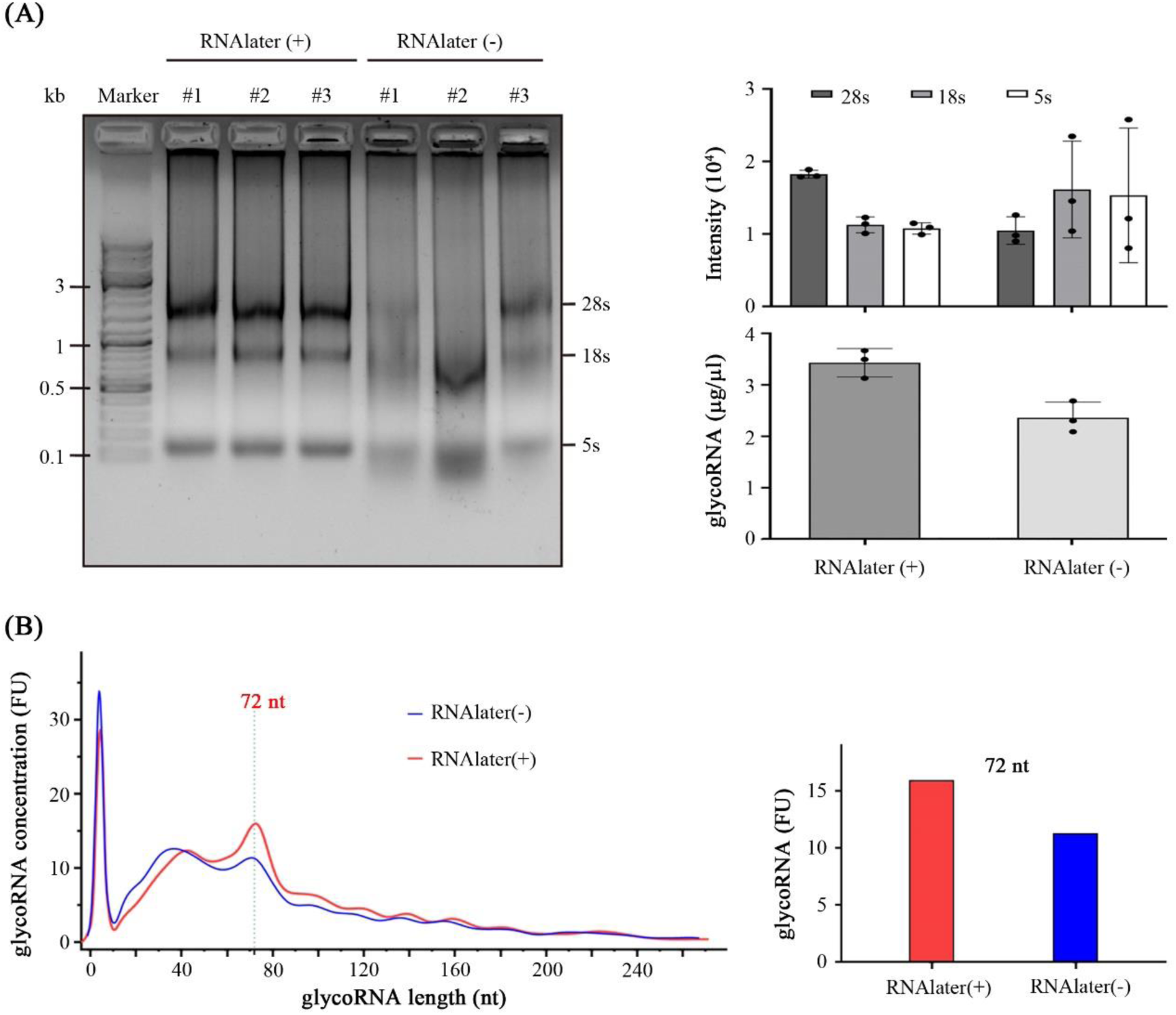
Improved RNA extraction using RNA protection buffer. (A) Gel electrophoresis of RNA isolated with and without RNAlater from tissues. Quantification of 28S, 18S, and 5S rRNA bands shows that RNAlater effectively preserves 28S and 18S rRNA, while 28S rRNA is significantly degraded without RNAlater protection. Additionally, the amount of glycoRNAs is much higher when tissue is stored in RNAlater buffer. (B) Comparison of glycoRNA concentration in tissues stored in RNAlater buffer. Longer glycoRNAs are observed in RNAlater-protected buffer, with the 72-nucleotide length (nt) band showing a higher concentration of glycoRNAs in RNAlater-treated tissues.

The Agilent Small RNA Kit for the 2100 Bioanalyzer Systems was used to analyze glycoRNA enrichment. This analysis generated multiple datasets with nucleotide length (nt) on the x-axis and fluorescence signal (FU) on the y-axis. To create an average representation, the “Average Multiple Curves” function within Origin 2022’s analysis-mathematics tools was employed. Subsequently, smoothing was applied to the averaged data to generate the final trend curve shown in **Figure 2**B. The results demonstrate that glycoRNAs from RNAlater-preserved tissues exhibit both higher concentrations and longer nucleotide lengths compared to those from empty tubes. Notably, the abundance of 72 nt glycoRNA is significantly higher in RNAlater-preserved tissues. These findings indicate that RNAlater not only prevents RNA degradation but also preserves the integrity of glycoRNAs.

### Impact of GAO oxidation on glycoRNA enrichment

To optimize glycoRNA enrichment using SPCgRNA or TnORNA, we assessed the effects of galactose oxidase (GAO) oxidation duration and enzyme units. We varied the GAO incubation time from 0.5 to 3 hours while keeping the initial total RNA amount constant. The amount of GAO units tested ranged from 0.5 U to 3 U. Gel electrophoresis analysis revealed that the signal intensity for oxidized RNA peaked at 2 hours (**Figure 3**A), suggesting optimal oxidation of most glycoRNAs at this time point. Interestingly, increasing GAO units did not always correlate with higher glycoRNA enrichment (**Figure 3**B). This could be attributed to potential RNA degradation or damage by excess GAO, leading to lower yields. Our findings emphasize the importance of balancing oxidation time and GAO units for optimal efficiency while preserving RNA integrity. We identified a reaction time of 2 hours with 1 U of GAO as the most effective for glycoRNA enrichment under these specific conditions.

**Figure 3.**
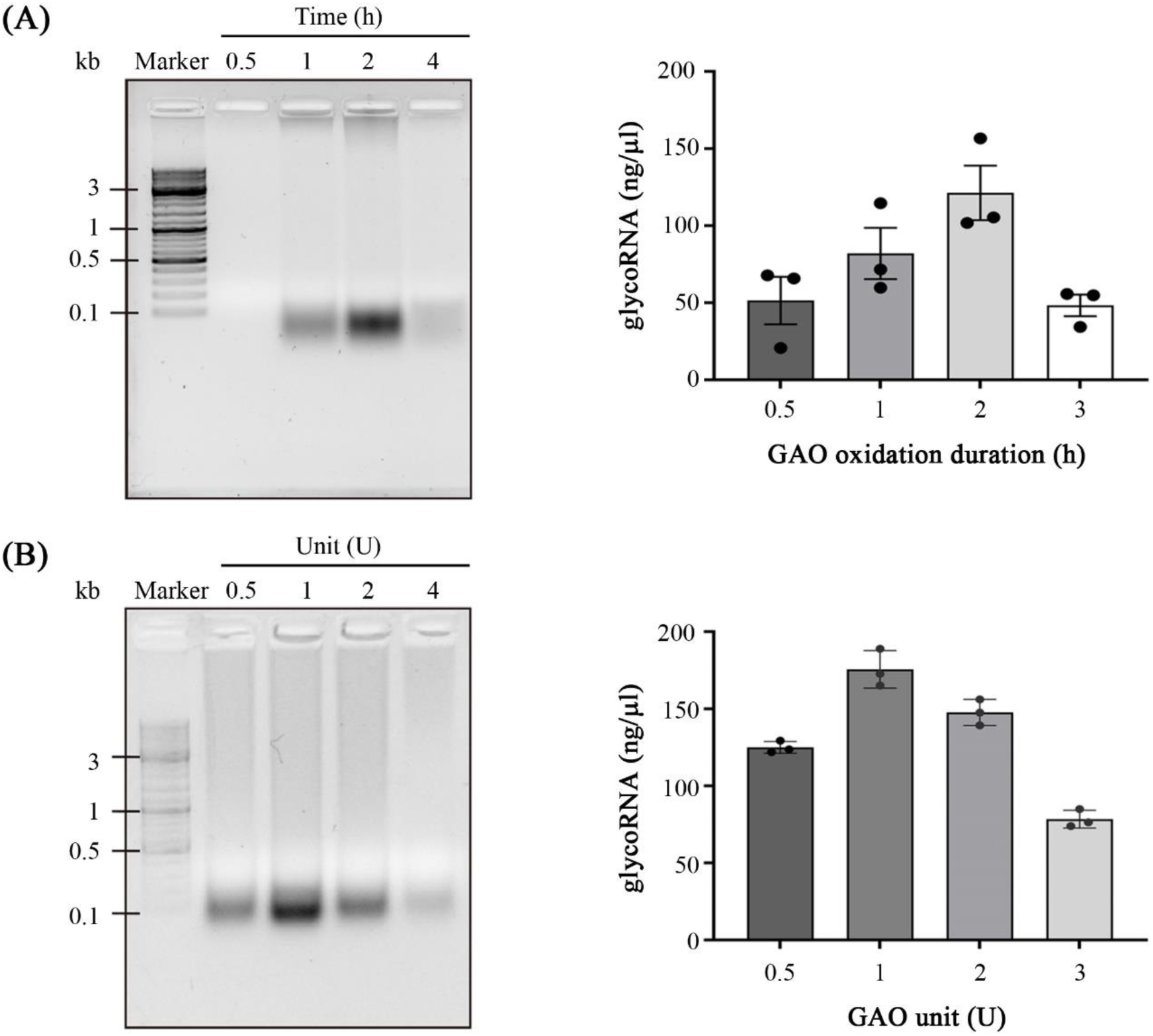
The effect of oxidation duration and amount of galactose oxidase on glycoRNA enrichment. (A) Duration of conjugation reaction (0.5, 1.0, 2.0, and 4.0 hours). (B) The amount of galactose oxidase (GAO) used for RNA oxidation.

### Optimizing glycosidase digestion for glycoRNA enrichment

We investigated key parameters influencing optimal conditions for glycosidase digestion of glycoRNAs: digestion temperature, incubation time, buffer pH, and the effect of sequential two-enzyme digestion (**Figure 4**). Releasing conjugated glycoRNAs from the hydrazide resin at different temperatures (4°C, 25°C, and 37°C) revealed a slightly higher yield at 25°C (**Figure 4**A). This might be due to a balance between efficient PNGase F digestion of N-glycoRNAs at moderate temperatures and reduced RNA degradation at lower temperatures. The trend curve also suggests that 37°C digestion results in shorter glycoRNA fragments compared to 25°C. Similarly, extending the incubation time from 1.5 hours to 3 hours led to a slight increase in glycoRNA concentration (**Figure 4**B). Buffer pH significantly impacted digestion efficiency. The highest glycoRNA concentration was observed after digestion in NH4HCO3 buffer (pH 8.0), likely due to optimal PNGase F activity at this pH (**Figure 4**C).

**Figure 4.**
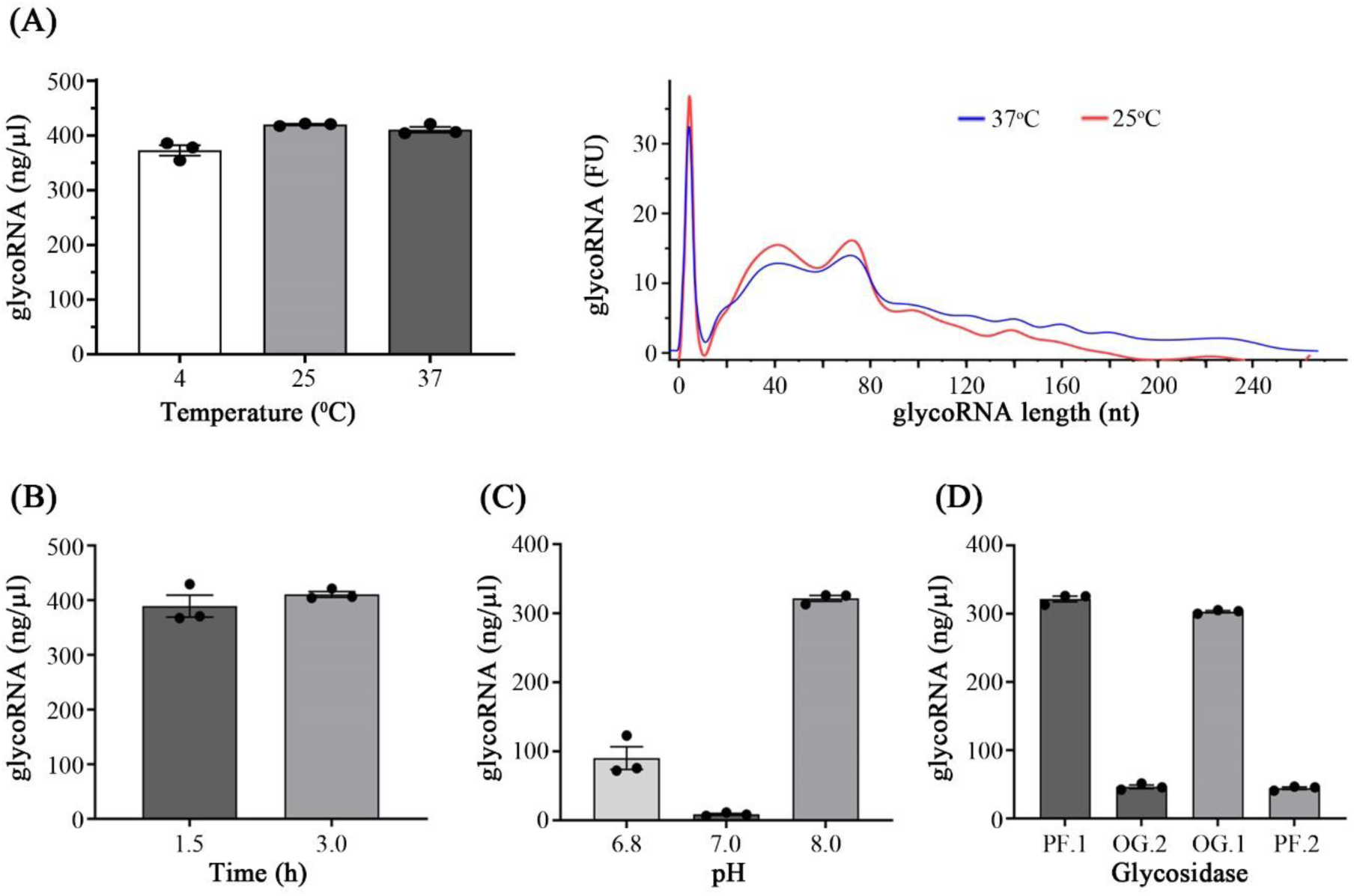
Effect of enzymatic digestion on glycoRNA enrichment. (A) Amount of glycoRNA obtained by PNGase F digestion at different temperatures (4/25/37°C). The distribution of glycoRNAs ranges from 0 to 260 nt, with the most abundant species being between 20 and 120 nt. (B) Effect of PNGase F digestion time. (C) Effect of different pH conditions on PNGase F digestion. The tested conditions include 25 mM Tris.HCl (pH 6.8), DEPC water (pH 7.0), and 25 mM ammonium bicarbonate (pH 8.0). (D) Sequential digestion of glycoRNA using a combination of glycosidases. PF.1 denotes first digestion with PNGase F, OG.2 denotes sequential digestion with a mixture of O-glycosidase and GalNAcEXO following PNGase F, OG.1 denotes first digestion with the O-glycosidase and GalNAcEXO mixture, and PF.2 denotes sequential digestion with PNGase F following the mixture.

We explored the possibility of simultaneously cleaving both N-glycoRNAs and O-glycoRNAs from a single sample using a sequential two-enzyme digestion on hydrazide resin. This approach involved PNGase F and a combination of O-glycosidase and GalNAcEXO. We tested two different enzyme orders: cleaving N-glycoRNAs first with PNGase F, followed by O-glycoRNAs with the combined enzymes, and vice versa. Our results revealed a significant decrease in glycoRNA concentration after the second enzyme digestion compared to the first, regardless of the order (**Figure 4**D). This suggests potential RNA degradation during the initial enzyme digestion step, even though the same sample was used. Therefore, for optimal enrichment of specific glycoRNA types, we recommend separate enzymatic cleavage approaches.

## Conclusion

This protocol outlines the optimal conditions for SPCgRNA or TnORNA methods to enrich and identify glycoRNAs. To ensure successful analysis, follow these steps: First, freshly collected tissues are immediately stored in an RNase-free RNA stabilization buffer, such as RNAlater, RNAsafe, or Monarch RNA Protection Reagent (NEB), to prevent RNA degradation. Next, RNA oxidation is performed by treating the total RNA with 1-2 U of freshly prepared galactose oxidase (GAO) for 2 hours. This crucial step helps maintain glycoRNA integrity while preventing over-oxidation. Finally, enzymatic digestion is conducted at room temperature. The appropriate enzyme should be selected according to the target RNA glycosylation, e.g., PNGase F for N-glycans or O-glycosidase/GalNAcEXO for O-glycans. Sample is usually incubated 1.5- to 3-hour in 25 mM NH4HCO3 (pH 8.0) for optimal cleavage of the glycoRNAs with minimal RNA degradation. While longer incubations (up to 3 hours) may improve enrichment, carefully monitor for RNA degradation to ensure successful analysis.

This technique applies to studying glycoRNAs in tissues and human body fluids, which can be isolated using various tools like vacuum-assisted filtration ^24^. These non-coding, single-stranded RNAs are present in blood, saliva, urine, and breast milk, making them potentially valuable disease markers ^25^. Notably, small RNAs are considered promising therapeutic targets for various diseases, including tumor-associated miRNAs ^26^ and rRNA-based antibiotics ^27^. Therefore, our method can be used to investigate whether abnormal glycosylation patterns on glycoRNAs can serve as diagnostic tools for disease onset and progression.

## Acknowledgments

This research was financially supported by the Soochow University Start-up Fund. We are grateful for additional funding from the Priority Academic Program Development of Jiangsu Higher Education Institutes (PAPD), the Jiangsu Science and Technology Plan Funding (BX2022023), the Jiangsu Shuangchuang Boshi Funding (JSSCBS20210697), the Suzhou Medical Innovation Funding (SKJY2021141), and the Suzhou Health Talents program (#9) (GSWS2022087).

## Author Contributions

X.Y.M. and J.B.D. conducted experiments, drafted the experimental sections, and analyzed data. X.T.W., S.Y.W., and Z.Y.Z. collected clinical samples and prepared samples. S.Y. drafted the manuscript. X.T.W., X.D.Y., and S.Y. provided funding support. X.D.Y. and S.Y. revised the manuscript. All authors approved this manuscript.

## Graphic Abstract

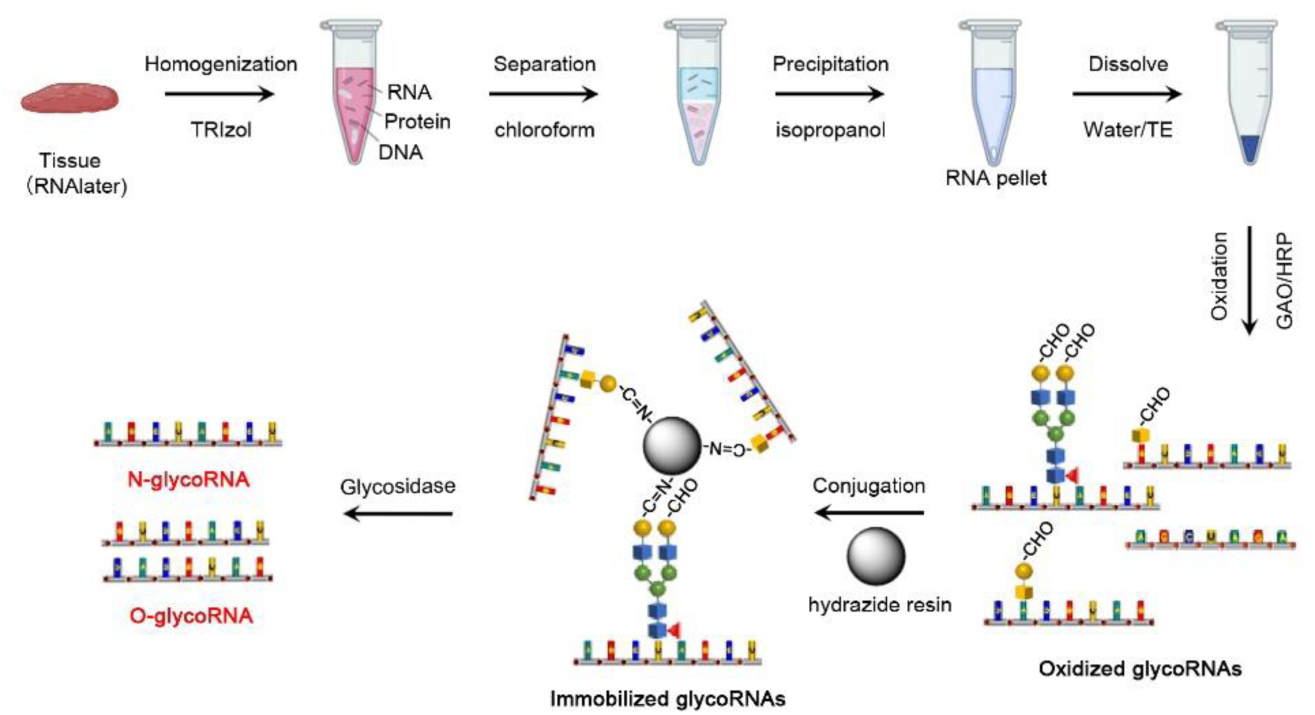

